# Oral immunization with a probiotic cholera vaccine induces broad protective immunity against *Vibrio cholerae* colonization and disease in mice

**DOI:** 10.1101/554303

**Authors:** Brandon Sit, Ting Zhang, Bolutife Fakoya, Aklima Akter, Rajib Biswas, Edward T. Ryan, Matthew K. Waldor

## Abstract

Oral cholera vaccines (OCVs) are being increasingly employed, but current killed formulations generally require multiple doses and lack efficacy in young children. We recently developed a new live-attenuated OCV candidate (HaitiV) derived from a *Vibrio cholerae* strain isolated during the 2010 Haiti cholera epidemic. HaitiV exhibited an unexpected probiotic-like activity in infant rabbits, preventing intestinal colonization and disease by wild-type *V. cholerae* before the onset of adaptive immunity. However, it remained unknown whether HaitiV would behave similarly to other OCVs to stimulate adaptive immunity against *V. cholerae.* Here, we orally immunized adult germ-free female mice to test HaitiV’s immunogenicity. HaitiV safely and stably colonized vaccinated mice and induced known adaptive immune correlates of cholera protection within 14 days of administration. Pups born to immunized mice were protected against lethal challenges of both homologous and heterologous *V. cholerae* strains. Cross-fostering experiments revealed that protection was not dependent on vaccine colonization in or transmission to the pups. These findings demonstrate the protective immunogenicity of HaitiV and support its development as a new tool for limiting cholera.

**Author summary:** Vaccines for cholera are gaining acceptance as public health tools for prevention of cholera and curtailing the spread of outbreaks. However, current killed vaccines provide minimal protection in young children, who are especially susceptible to this diarrheal disease, and do not stimulate immunity against antigens that may only be expressed by live bacteria during infection. We recently developed HaitiV, an extensively engineered live-attenuated oral cholera vaccine candidate, derived from a clinical isolate from the Haiti cholera outbreak. Here, we found that the HaitiV induces immunological correlates of protection against cholera in germ free mice and leads to protection against disease in their offspring. Protection in this model was dependent on passively acquired factors in the milk of immunized dams and not transmission or colonization of HaitiV. Coupling the immunogenicity data presented here with our previous observation that HaitiV can protect from cholera prior to the induction of adaptive immunity, suggests that HaitiV may provide both rapid-onset short-term protection from disease while eliciting stable and long-lasting immunity against cholera.

## Introduction

The bacterial pathogen *Vibrio cholerae* causes the severe human diarrheal disease cholera, a potentially fatal illness characterized by rapid-onset of fluid loss and dehydration. Recent estimates place the global burden of cholera at ~3 million cases per year, and over 1.3 billion people are at risk of this disease [1]. *V. cholerae* proliferates in the small intestine and produces cholera toxin (CT), which leads to water and electrolyte secretion into the intestinal lumen [2].

The O1 serogroup of *V. cholerae* causes virtually all epidemic cholera. This serogroup includes two serotypes, Inaba and Ogawa, whose LPS structures differ by a single methyl group on the terminal O-antigen sugar [3]. Serologic and epidemiologic studies have established the existence of extensive serotype cross-reactivity and -protectivity, although immunogenicity and protection is highest to the homologous serotype [4–7]. Toxigenic O1 strains are divided into two major biotypes, classical and El Tor, but the former has not been isolated in over a decade and is thought to be extinct [8]. Ongoing evolution of El Tor *V. cholerae* has given rise to “atypical El Tor” (AET) strains, which are distinguishable from earlier strains by a variety of features, including the expression of a non-canonical *ctxB* allele, that may impact disease severity in afflicted patients [4,9,10]. AET strains, such as the strain responsible for the 2010 Haitian cholera epidemic, are thought to be the globally dominant cause of cholera [10–12]. Currently, serogroup O139 isolates only cause sporadic disease [13]. Notably, antibodies (or immune responses) targeting the O1 O-antigen do not protect against O139 challenge and vice versa [14–16].

Oral cholera vaccines (OCVs) have recently become widely accepted as a tool for cholera control [17]. Vaccines are a potent method to combat cholera due to their ability to both directly and indirectly reduce disease and transmission [18]. Killed multivalent whole-cell OCVs, such as Shancol, have shown promise both to prevent disease in endemic regions and as reactive agents to limit cholera during epidemics [19]. However, killed OCVs tend to be less effective at eliciting protective immunity in young children (<5 years old), who are most susceptible to cholera [20,21]. Additionally, these vaccines typically require two doses over the span of several weeks, although recent studies suggest that a single dose may still lead to moderate protection [20,22,23].

There is no live-attenuated OCV licensed for use in cholera-endemic regions. The only clinically available live-attenuated OCV is Vaxchora (CVD103-HgR), which is derived from a classical O1 Inaba *V. cholerae* strain and was approved by the US FDA in 2017 for use in travelers [24]. In contrast to killed OCVs, live vaccines, such as CVD103-HgR and the El Tor-derived vaccine Peru-15, elicit more potent immune responses in young children [25,26], potentially because they more closely mimic natural infection than killed OCVs. In particular, live vaccines can produce antigens in vivo that are not expressed in the in vitro growth conditions used to prepare killed vaccines; furthermore, the inactivation processes used to formulate killed vaccines can destroy antigenic epitopes [27].

In addition to the requirement for multiple doses of some OCVs for optimal protection, all current live and killed OCVs are thought to be accompanied by a post-vaccination lag in protection during induction of anti-*V. cholerae* adaptive immunity. The shortest reported time to protective efficacy is 8 days post-vaccination, a delay that could hamper reactive vaccination campaigns designed to limit the spread of cholera outbreaks [28]. We recently created HaitiV, a new live-attenuated OCV candidate derived from an AET O1 Ogawa *V. cholerae* clinical isolate from the 2010 Haiti cholera outbreak. HaitiV harbors many genetic alterations that render it avirulent and resistant to reversion while preserving its robust capacity for colonization of the small intestine [29]. In an infant rabbit model of cholera [30], intestinal colonization with HaitiV conferred protection against lethal wild-type (WT) *V. cholerae* challenge within 24 hours of vaccination, a timescale inconsistent with the development of adaptive immunity and suggestive of a “probiotic”-like mechanism of protection. Here, using a mouse model of *V. cholerae* intestinal colonization, we show that oral administration of HaitiV to female mice elicits serum vibriocidal antibodies and protects their pups from lethal challenge with virulent *V. cholerae*. Thus, HaitiV has the potential to provide rapid probiotic-like protection as well as to elicit long-lasting immune protection from cholera.

## Methods

### Bacterial strains and growth conditions

All bacteria were grown in Luria-Bertani (LB) broth supplemented with the relevant chemicals at the following concentrations: streptomycin (Sm, 200μg/mL), kanamycin (200μg/mL), carbenicillin (Cb, 50μg/mL), sulfamethoxazole/trimethoprim (SXT, 80 and 16μg/mL) and 5-bromo-4-chloro-3-indolyl-β-d-galactopyranoside (X-gal, 60μg/mL). For growth on plates, LB + 1.5% agar was used. All *V. cholerae* strains in this study were spontaneous Sm^R^ derivatives of the wild-type. Bacteria were stored as −80°C stocks in LB with 35% glycerol. A list of strains and plasmids used in this study is listed in Supplementary Table 1.

### Construction of *ΔctxAB* Haiti *V. cholerae*

The CT deletion strain in the H1 *V. cholerae* background (HaitiWT) was generated by allelic exchange as previously described, with an additional selection step to enhance the efficiency of obtaining a stable single crossover strain [29]. Briefly, HaitiWT was conjugated with SM10pir *E. coli* bearing the suicide plasmid pCVD442-ctxAB-KnR, containing *sacB* as well as a kanamycin resistance cassette from pKD4 sandwiched by homology arms targeting the *ctxAB* operon (locus tags N900_RS07040 – N900_RS07045). Single crossovers were selected on LB+Sm/Cb/Kn agar plates. To select for a double crossover, verified single crossovers were grown in LB + Cb/Kn for 4 hours at 37°C and then passaged in LB+10% sucrose overnight at room temperature. Sucrose-resistant (*sacB*-negative), Kn^R^ and Cb^S^ colonies were then conjugated with SM10pir *E. coli* bearing pCVD442-ctxAB (no Kn^R^ cassette) and clean Kn^S^ double crossovers generated via an identical protocol. The *ΔctxAB* deletion was verified by colony PCR with internal and flanking primers.

### Oral immunization regimen

4-week old germ-free (GF) female C57BL/6 (Massachusetts Host-Microbiome Center) or Swiss-Webster (Taconic Farms) mice were housed in a BL-2 facility for the duration of the experiment. On Day 0, 2, 4, 6, 14, 28, 42 and 56, mice were anesthetized with isoflurane and orally gavaged with 10^9^ CFU of an overnight culture of either HaitiV or CVD103-HgR in 100μL 2.5% Na_2_CO_3_. Mice were weighed at every immunization and once every 4-5 days between boosts. At each weighing, fresh fecal pellets were plated on LB + Sm to enumerate shed bacteria. At Day 7, 14, 28 and 42 post immunization, blood samples were obtained from each mouse by tail vein incision. A Day 1 blood sample was collected from the Swiss-Webster cohort and the single-dose C57BL/6 cohort. Blood was clotted at room temperature for 1 hour, centrifuged at 20000 x g for 5 minutes and the supernatant (serum) stored at −20°C for analysis.

### Quantification of vibriocidal responses

Vibriocidal antibody quantification was performed by complement-mediated cell lysis using PIC018 (Inaba) or PIC158 (Ogawa) *V. cholerae* as the target strain as previously described [31]. Seroconversion was defined as ≥4x increase in titer over the baseline measurement. The characterized mouse monoclonal antibody 432A.1G8.G1.H12 targeting *V. cholerae* O1 OSP was used as a positive control for the vibriocidal assay. Titers are reported as the dilution of serum causing a 50% reduction in target optical density compared to no serum control wells.

### Quantification of anti-CtxB and anti-OSP responses

Anti-CtxB and anti-OSP responses were measured by previously described isotype-specific ELISAs [32,33]. Briefly, 96-well plates (Nunc) were coated with 1 μg/mL solution of bovine GM1 monosialoganglioside (Sigma) in 50mM carbonate buffer overnight. Next, 1μg/mL CtxB in 0.1% BSA/PBS purified from the classical Inaba strain 569B (List Biological Laboratories) was layered onto the GM1-coated wells. Wells were blocked with a 1% BSA/PBS mixture after which 1:50 dilutions of the mouse serum samples were loaded into each well. Goat anti-mouse IgA, IgG or IgM secondaries conjugated to HRP (Southern Biotechnology) were then added at a concentration of 1μg/mL in 0.1% BSA/0.05% Tween/PBS and incubated for 90 minutes. Detection was performed by adding an ABTS/H_2_O_2_ mixture to the wells and taking an absorbance measurement at 405nm with a Vmax microplate kinetic reader (Molecular Devices Corp., Sunnyvale, CA). Plates were read for 5 min at 30 s intervals, and the maximum slope for an optical density change of 0.2 U was reported as millioptical density units per minute (mOD/min). Results were normalized using pooled control serum from mice previously immunized against cholera and reported as ELISA Units as previously described [32]. Anti-OSP responses were measured and reported similarly to anti-CtxB responses, only instead of CtxB, purified OSP:BSA from either PIC018 or PIC158 (1 μg/mL) was used to coat plates as previously described [34]. Additionally, OSP ELISAs were carried out with 1:25 dilutions of the serum samples.

### Infant mouse challenge assay

The infant mouse survival challenge was adapted from previous reports to optimize the dosage for HaitiWT and to include more frequent monitoring intervals [31,32]. Pregnant dams were singly housed at E18-19 for delivery. At P3 (third day of life), pups were orally inoculated with 10^7^ CFU *V. cholerae* in 50μL LB and returned to their dam. Infected pups were monitored every 4-6 hours for onset of diarrhea and reduced body temperature. Once symptoms appeared, monitoring was increased to every 30 minutes until moribundity was reached, at which point pups were removed from the nest and euthanized by isoflurane inhalation followed by decapitation for dissection and CFU plating of the small intestine on LB + Sm/X-gal. Pups that were alive at 48 hpi were deemed protected from the challenge. Cross-fostering was performed by transferring up to half of a litter between dams on the first day of life (P1). Fostering was maintained for at least 48 hours before infection to fully replace the milk from the original dam. We excluded rejected pups from analyses due to our inability to attribute mortality to infection alone.

### Statistical analysis

Statistical analyses were performed with Prism 8 (Graphpad). Due to missing values from paired measurements as a result of insufficient serum sampling, antibody titers could not be analyzed by a typical one-way repeated measures ANOVA. Instead, we employed a mixed-effect model ANOVA using the earliest sample (Day 1 or Day 7) as the control and performed post hoc tests with Dunnett’s multiple comparison test. The Geisser-Greenhouse correction was applied for the ANOVA. Survival curves were analyzed with the log-rank test and CFU burdens were compared with the Mann Whitney U test. A p-value <0.05 was considered statistically significant.

### Animal use statement

This study was performed in accordance with the NIH Guide for Use and Care of Laboratory animals and was approved by the Brigham and Women’s Hospital IACUC (Protocol #2016N000416). Infant (P14 or younger) mice were euthanized by isoflurane inhalation followed by decapitation. At the end of the study, adult mice were euthanized by isoflurane inhalation followed by cervical dislocation.

## Results

### Experimental design and vaccine colonization

While the infant rabbit model enables investigation of the progression of a *V. cholerae*-induced diarrheal disease that closely mimics human cholera [30], it is not appropriate to study vaccine immunogenicity because newborn animals lack a fully developed immune system. Instead, we used adult GF mice to study HaitiV immunogenicity. In contrast to normal adult mice, which are resistant to *V. cholerae* intestinal colonization, oral inoculation of GF mice with *V. cholerae* results in stable intestinal colonization without adverse effects [35–37]. In the GF model, serum markers of immunity, such as vibriocidal titers, can be measured, but challenge studies are not possible due to the persistent colonization of the vaccine strain and the resistance of adult mice to diarrheal disease. Here, we further developed a variation of the GF model [38]. Besides measuring serum markers in the orally vaccinated adult mice, neonatal pups (which are sensitive to *V. cholerae* induced diarrheal disease) born to these mice were subjected to challenge studies to evaluate vaccine protective efficacy.

We established two cohorts of orally immunized adult female GF mice. In the first cohort, a small pilot study was set up to compare the immunogenicity of HaitiV and a streptomycin-resistant derivative of CVD-103HgR. This cohort consisted of 4-week-old Swiss-Webster GF mice that were immunized with either vaccine strain (n=3 per group). Cohort 2 consisted of a set of seven 4-week-old C57BL/6 mice that were all immunized with HaitiV. We generally followed the multi-dose oral immunization scheme previously used in this model, which included eight doses of 1×10^9^ CFU vaccine over eight weeks [36,37]. After this vaccination regimen, the mice in cohort 2 were mated and vaccine-induced protective immunity was assessed in the progeny (Fig 1A).

**Fig 1.**
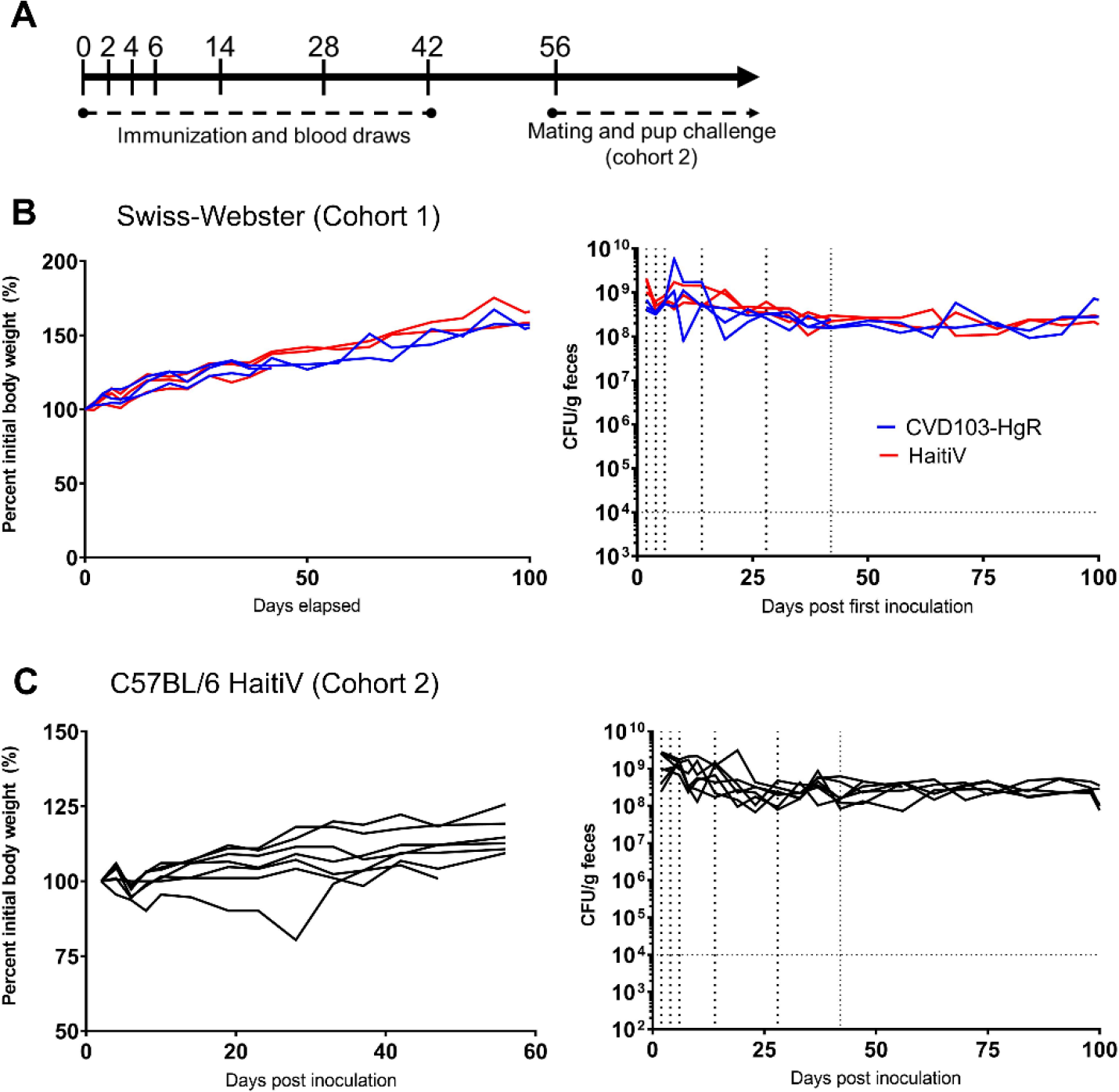
Fecal vaccine shedding and weight of adult GF mice after oral administration of live attenuated cholera vaccines. (A): Schematic of vaccination regime – note cohort 1 does not extend past Day 42. (B and C): Bodyweight (left) and fecal CFU shedding of HaitiV (right) for cohort 1 and 2 respectively.

Based on fecal CFU, all animals in both cohorts were stably colonized with high levels of either vaccine strain (Fig 1B). No adverse effects of long-term colonization with HaitiV or CVD103-HgR were noted, and all mice gained weight over the course of the study (Fig 1B). Fecal shedding and presumably intestinal colonization of HaitiV in cohort 2 was eliminated after these dams were used to cross-foster pups born to specified-pathogen free (SPF) control mice (described below), suggesting that a normal microbiota can outcompete HaitiV.

### HaitiV immunization elicits robust serum antibodies targeting *V. cholerae*

Serum samples from the immunized mice were used to quantify antibodies targeting several *V. cholerae* factors thought to play roles in protection from cholera. One of these metrics, the vibriocidal antibody titer, is a validated correlate of protection in vaccinated humans [39–42]. In cohort 1, all mice immunized with HaitiV or CVD-103HgR seroconverted within 2 weeks and developed vibriocidal titers consistent with those reported in human studies for live OCVs (Fig 2) [41,43]. Furthermore, HaitiV and CVD-103HgR elicited comparable vibriocidal titers.

**Fig 2.**
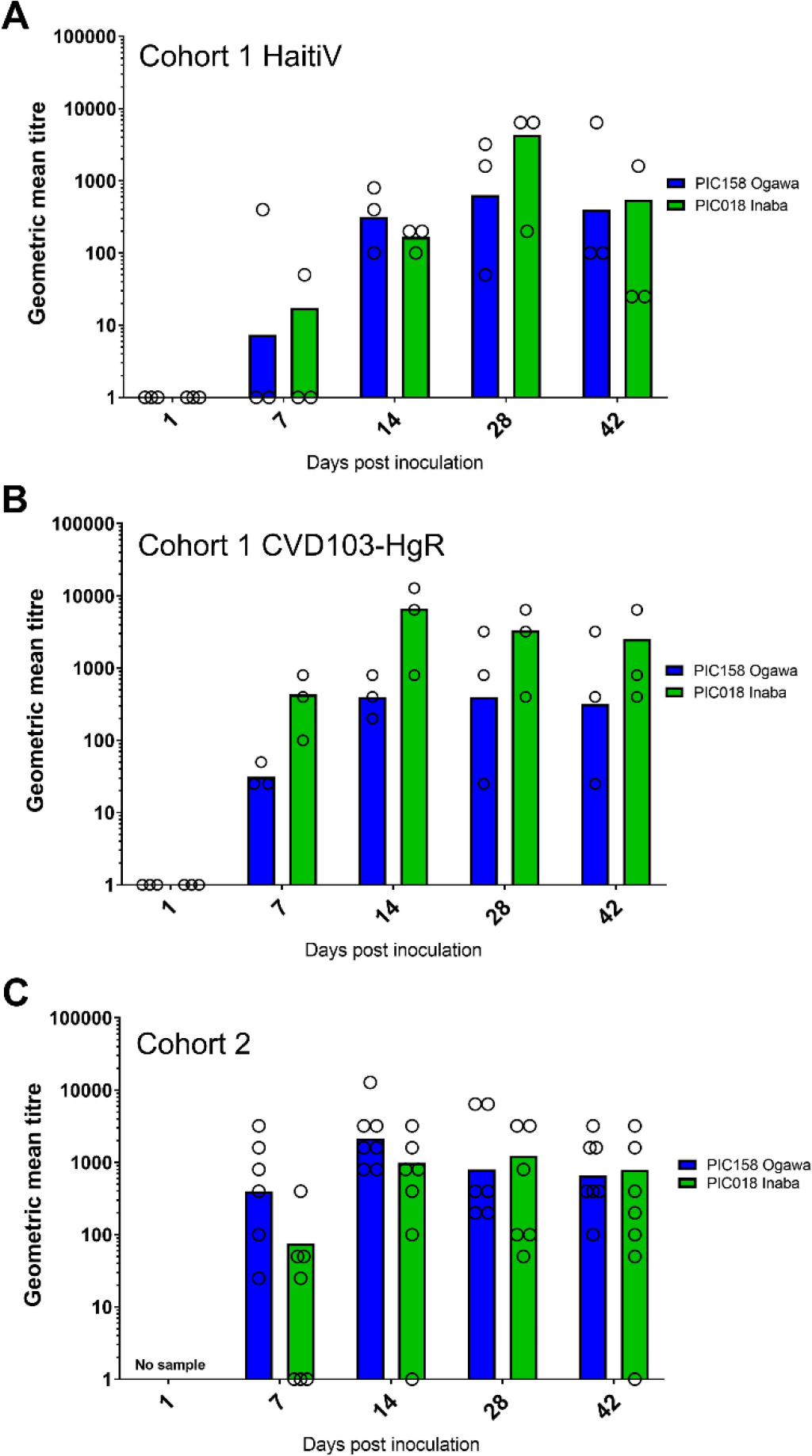
Vibriocidal titers in immunized mice. Vibriocidal titers measured in cohort 1 mice immunized with (A): HaitiV or (B): CVD103-HgR, and in (C): cohort 2 mice immunized with HaitiV are shown. Each circle corresponds to the lowest dilution at which vibriocidal activity was observed and the height of the bars represent geometric mean titers in each group. Ogawa-specific responses, measured using strain PIC158, are shown in blue and Inaba-specific responses, measured using strain PIC018, are shown in green. Serum samples with undetectable vibriocidal activity were assigned a titer of 1

In cohort 2, HaitiV immunization of C57BL/6 mice also induced high vibriocidal titers to Ogawa and Inaba target strains (Fig 2C). Isotype-specific levels of antibodies targeting Ogawa and Inaba O-antigen specific polysaccharide (OSP), and the B-subunit of CT (CtxB) were also measured since they also likely contribute to immunity to cholera [39]. Although we did not measure Day 1 titers in cohort 2, measurements from naïve GF C57BL/6 mice and baseline measurements from cohort 1, and Day 1 of HaitiV-inoculated C57BL/6 mice in a later cohort (Fig S2) showed undetectable levels of vibriocidal antibodies (Fig 2A, S2). The cohort 2 mice developed strong anti-Ogawa and anti-Inaba OSP responses (Fig 3, Table S2). The anti-Ogawa OSP titers were generally higher than those targeting Inaba OSP, likely reflecting the fact that HaitiV is an Ogawa strain. All mice in cohort 2 also developed high levels of anti-CtxB IgA, IgG and IgM antibodies (Fig 4, Table S1). The 100% seroconversion rate and general increase over time of all three humoral immune responses measured (vibriocidal, anti-CtxB and anti-OSP antibodies) reveals that orally delivered HaitiV can elicit *V. cholerae*-specific immune responses.

**Fig 3.**
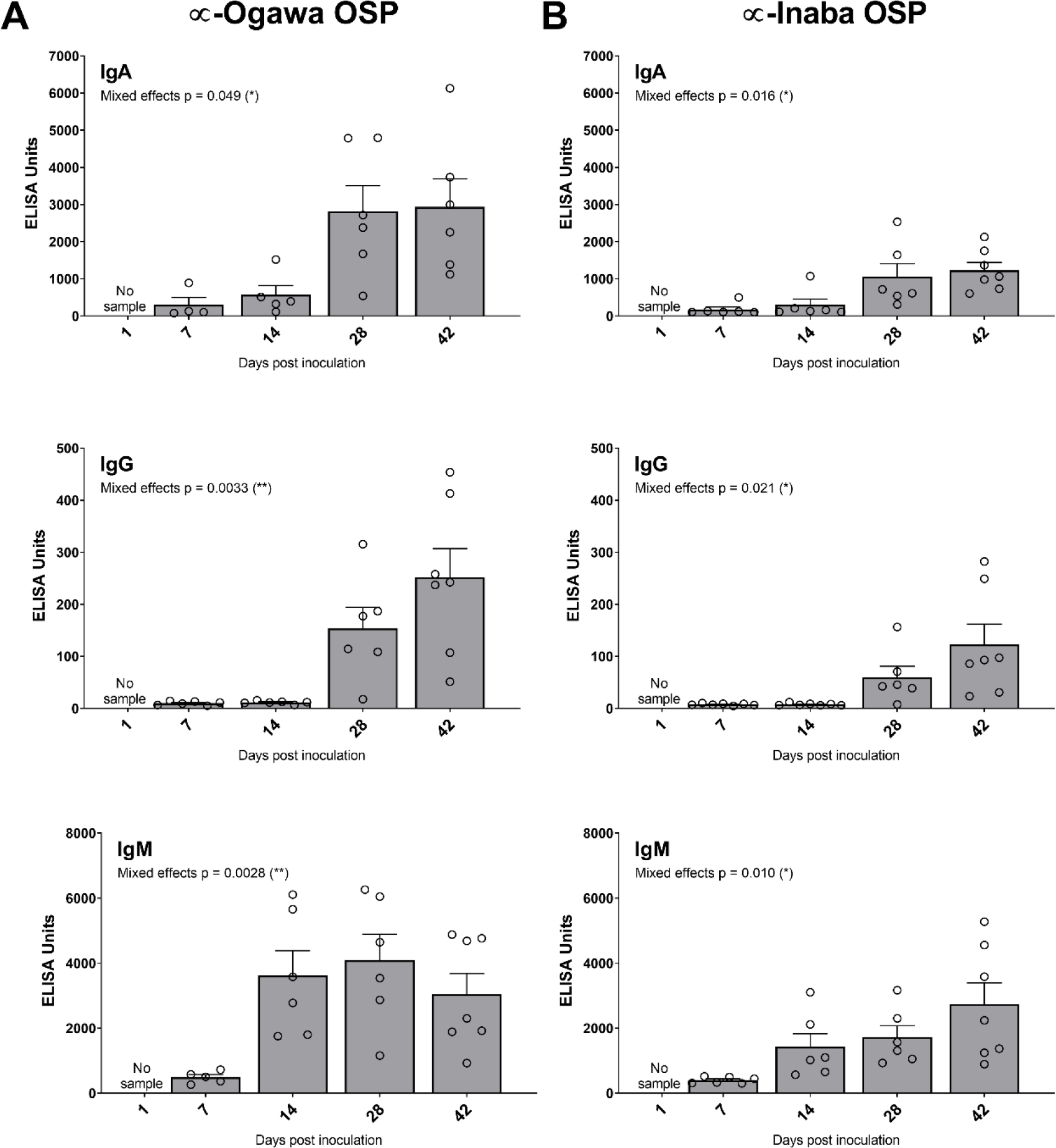
Ogawa- and Inaba-specific anti-OSP titers in C57BL/6 mice immunized with HaitiV. Anti-Ogawa OSP titers with specific isotype data for IgA (top), IgG (middle) and IgM (bottom). Anti-Ogawa OSP titers with specific isotype data for IgA (top), IgG (middle) and IgM (bottom). The mean ± SEM is indicated. Mixed-effect p-values were determined by one-way ANOVA analysis. Multiple comparison p-values for this data are reported in Table S1. Although there were 7 mice per group, some samples were of insufficient volume for analysis.

**Fig 4.**
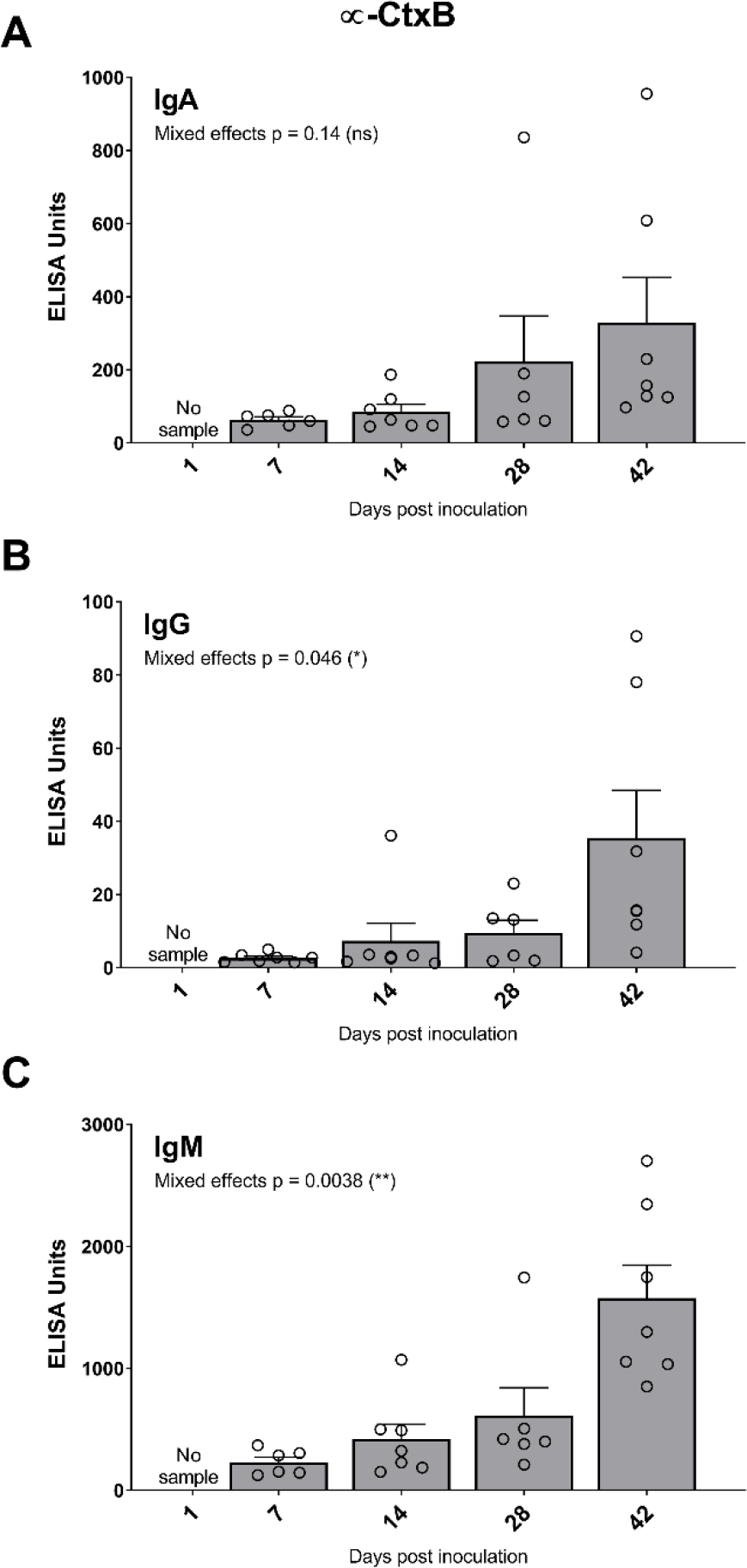
Isotype specific anti-CtxB titers in C57BL/6 mice immunized with HaitiV. Anti-CtxB titers with specific isotype analysis for IgA (A), IgG (B) and IgM (C) were performed for each serum sample. The mean ± SEM is indicated. Mixed-effect p-values were determined by one-way ANOVA analysis. Multiple comparison p-values for this data are reported in Table S1. Although there were 7 mice per group, some samples were of insufficient volume for analysis.

### Pups born to HaitiV immunized dams are protected from lethal *V. cholerae* challenge

To assess the protective efficacy of HaitiV in this model, we challenged the neonatal progeny of HaitiV-immunized or control dams with lethal doses of different wild type *V. cholerae* strains. This assay has been used to study passive immunity elicited by cholera vaccines, but has not been characterized in vaccinated GF mice [31,32]. Initially, we optimized this assay with litters from SPF C57BL/6 control mice. Three or four-day old pups were inoculated with 10^7^ or 10^8^ CFU of HaitiWT, the virulent strain from which HaitiV was derived and returned to their dams for monitoring (Fig 5A). Infected pups from both groups rapidly developed signs of dehydrating diarrheal disease, including accumulation of nest material on their anogenital regions, lethargy, skin tenting and hypothermia. All infected pups died by 48 hours post inoculation (hpi), with a median time to moribundity of ~23-26 hpi (Fig 5B). At the time of death, all pups were heavily colonized, with >10^7^ CFU/small intestine (Fig 5C), and had swollen ceca, another hallmark of productive cholera infection in mammalian models [30,44]. Since there were no significant differences in survival or bacterial loads in mice challenged with either 10^7^ or 10^8^ CFU, the smaller dose was used in subsequent experiments (Fig 5B). Diarrhea and death in this model were entirely dependent on CT; infant mice inoculated with HaitiWT Δ*ctxAB* or HaitiV were completely healthy at 48 hpi, despite sustained intestinal colonization (Fig 5C).

**Fig 5.**
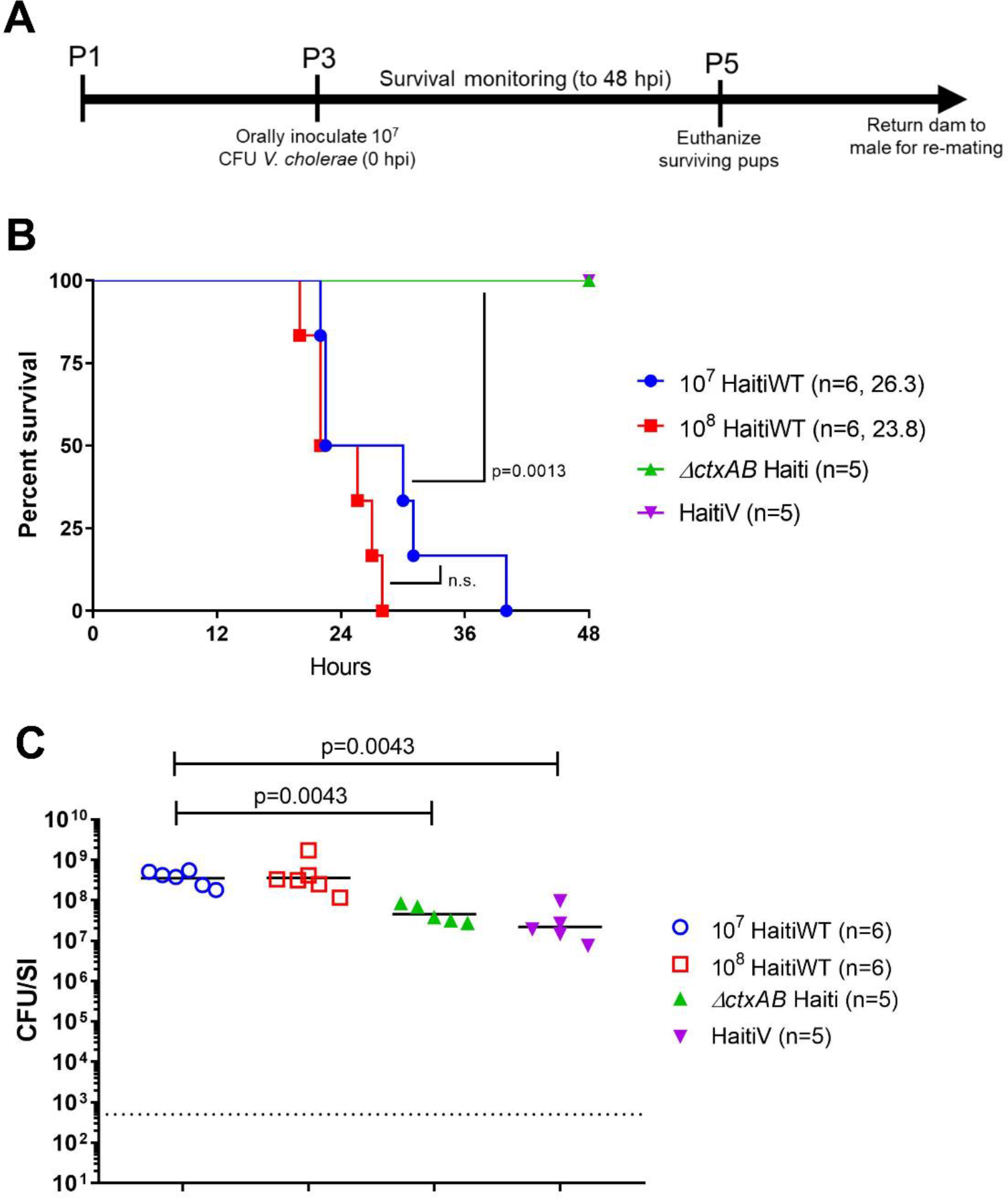
Survival of P3 C57BL/6 pups challenged with Haiti *V. cholerae*. (A): Timeline of oral challenge assays. (B) Survival curves for 48 hours post inoculation with the indicated strains; the number of pups challenged and the median survival time in hours is shown. P-values were determined by the Mantel-Cox method. (C) Intestinal *V. cholerae* burden at the time of death (open shapes) or at 48 hours (filled triangles). Burden is plotted as colony-forming units per small intestine (CFU/SI). The dotted line marks the limit of detection (50 CFU/SI). P-values were determined by the Mann-Whitney U test.

We next mated HaitiV-immunized animals from cohort 2 with age-matched GF male mice, thereby preserving their colonization with HaitiV. When challenged with HaitiWT, none of the 16 pups born to HaitiV-immunized dams developed signs of diarrhea or died by 48 hpi; in stark contrast, all pups born to non-immunized dams died within ~30 hpi (Fig 6A, left). There was a marked ~5,000-fold reduction in the intestinal load of HaitiWT in pups born to immunized versus control dams (Fig 6A, right). The pups of the immunized dams remained healthy for at least 2 weeks post-challenge, even though there were still detectable but very low levels of HaitiWT in their intestinal homogenates (Fig S1). Thus, oral immunization with HaitiV elicits an immune response that provides potent protection in nursing pups from diarrheal disease, death and *V. cholerae* intestinal colonization.

**Fig 6.**
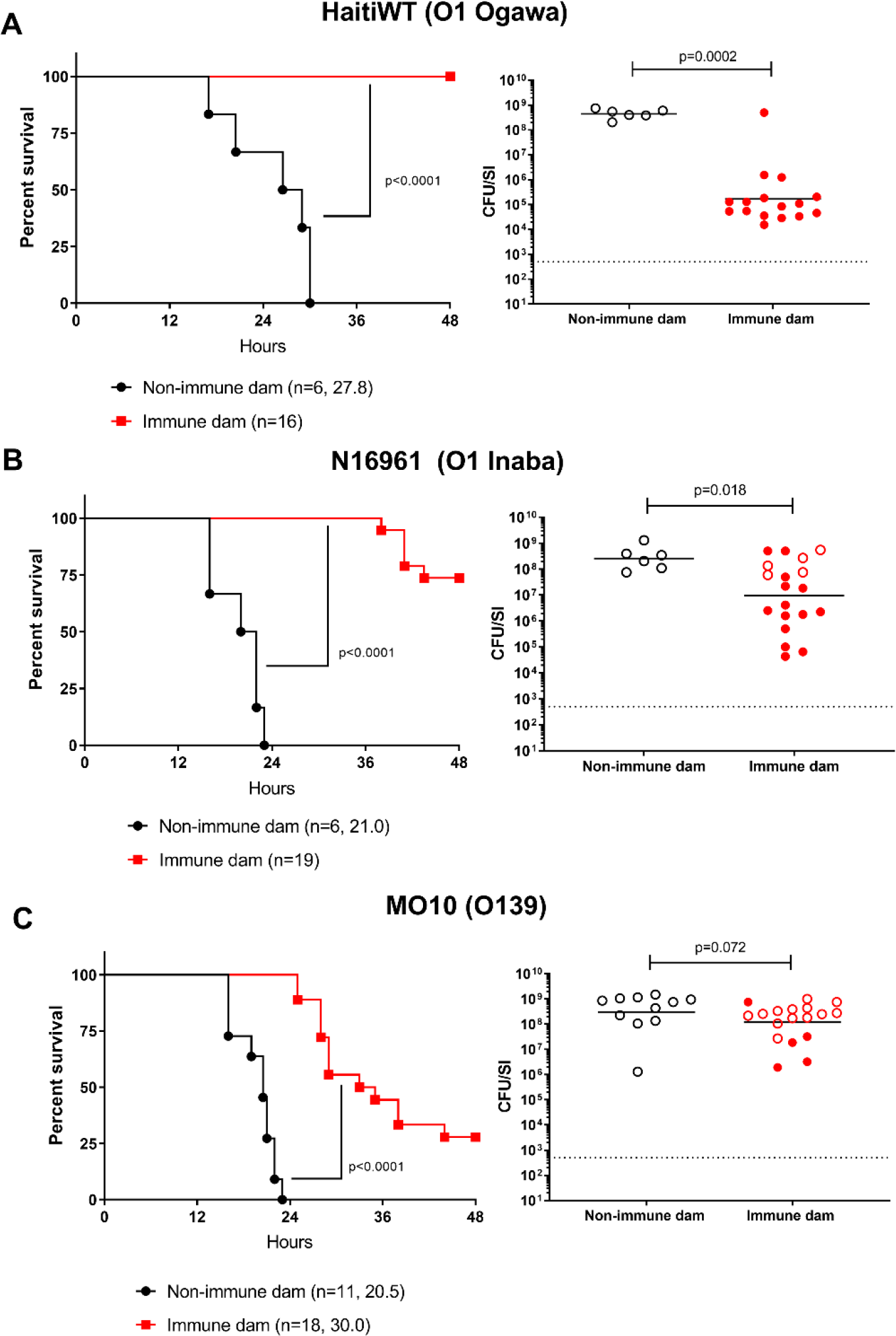
Survival of P3 C57BL/6 pups born to HaitiV-immunized or non-immunized dams after challenge with different serotype and serogroup *V. cholerae* strains. Pups born to either HaitiV-immunized dams (red) or non-immunized SPF dams (black) were challenged with either the O1 Ogawa strain HaitiWT (A), the O1 Inaba strain N16961 (B) or the O139 strain MO10 (C). Panels on the left show survival curves for infected pups up to 48 hours post inoculation with the indicated strains; the number of pups challenged and the median survival time in hours is shown. P-values were determined by the Mantel-Cox method. Panels on the right show the intestinal *V. cholerae* burden in pups at the time of death (open circles) or at 48 hpi (closed circles). Burden is plotted as CFU/SI. The dotted line marks the limit of detection (50 CFU/SI) and grey open circles indicate pups with burdens below this limit. P-values were determined by the Mann-Whitney U test.

Pups of HaitiV-immunized dams were similarly challenged with heterologous *V. cholerae* strains, to test the serotype and serogroup specificity of protection engendered by oral immunization with HaitiV. The additional challenge strains included an O1 Inaba strain (N16961) that has been used as the challenge strain in several human volunteer cholera studies [43,45] and the serogroup O139 strain MO10, which was isolated during the 1992 O139 outbreak in India. Most pups from HaitiV-immunized dams were protected from N16961 *V. cholerae* challenge (7/10 survival at 48 hpi, Fig 6B). Despite the clinical protection, there was a much less dramatic reduction in the intestinal burden of N16961 (~20-fold) compared to that observed with HaitiWT challenge, indicating serotype-specific responses play an important role in limiting colonization. Surprisingly, pups challenged with MO10 also exhibited some protection, but there was no concomitant reduction in the intestinal burden of this O139 strain (Fig 6C). Together, these observations demonstrate that animals can exhibit protection from death despite relatively robust colonization, suggesting that protection from disease may result from immunity targeting factors such as CtxB, in addition to those that impede colonization.

### Pups fostered by HaitiV-immunized dams are protected from lethal *V. cholerae* challenge

Since our earlier studies indicated that HaitiV itself can mediate rapid protection against cholera independent of an adaptive immune response, it was important to investigate whether pups nursed by HaitiV immunized dams were colonized with the vaccine strain. Extensive plating of intestinal samples from the >50 pups used for survival assays (limit of detection = 50 CFU/small bowel) did not reveal any HaitiV CFU in the pups reared by HaitiV-shedding dams. Thus, vaccine strain transmission and its probiotic effects are almost certainly not the explanation for the potent protection observed in nursing pups.

Cross-fostering experiments were undertaken to investigate the likely passive nature of the protection. P1 pups born to SPF dams were transferred to and reared by HaitiV-immunized dams and then challenged 2 days later with HaitiWT (Fig 5A, between P1-P3). All pups crossed-fostered by immunized dams were protected (100% survival at 48 hpi) and nearly all had marked reductions (~1,000 fold) in their intestinal HaitiWT burdens (Fig 7A). These observations mirror the challenge studies presented above (Fig 6A), indicating passive immunity from milk accounts for the protection that HaitiV-immunized dams bestow to their progeny. Conversely, when pups born to HaitiV-immunized dams were cross-fostered by SPF (non-vaccinated) dams, all succumbed to HaitiWT challenge, albeit with an increase in median survival time (by ~6-hour) and had high HaitiWT intestinal burdens (Fig 7B). The modest extension in survival time in these mice may be due to trans-placentally derived immunity or residual milk from the HaitiV-immunized dam.

**Fig 7.**
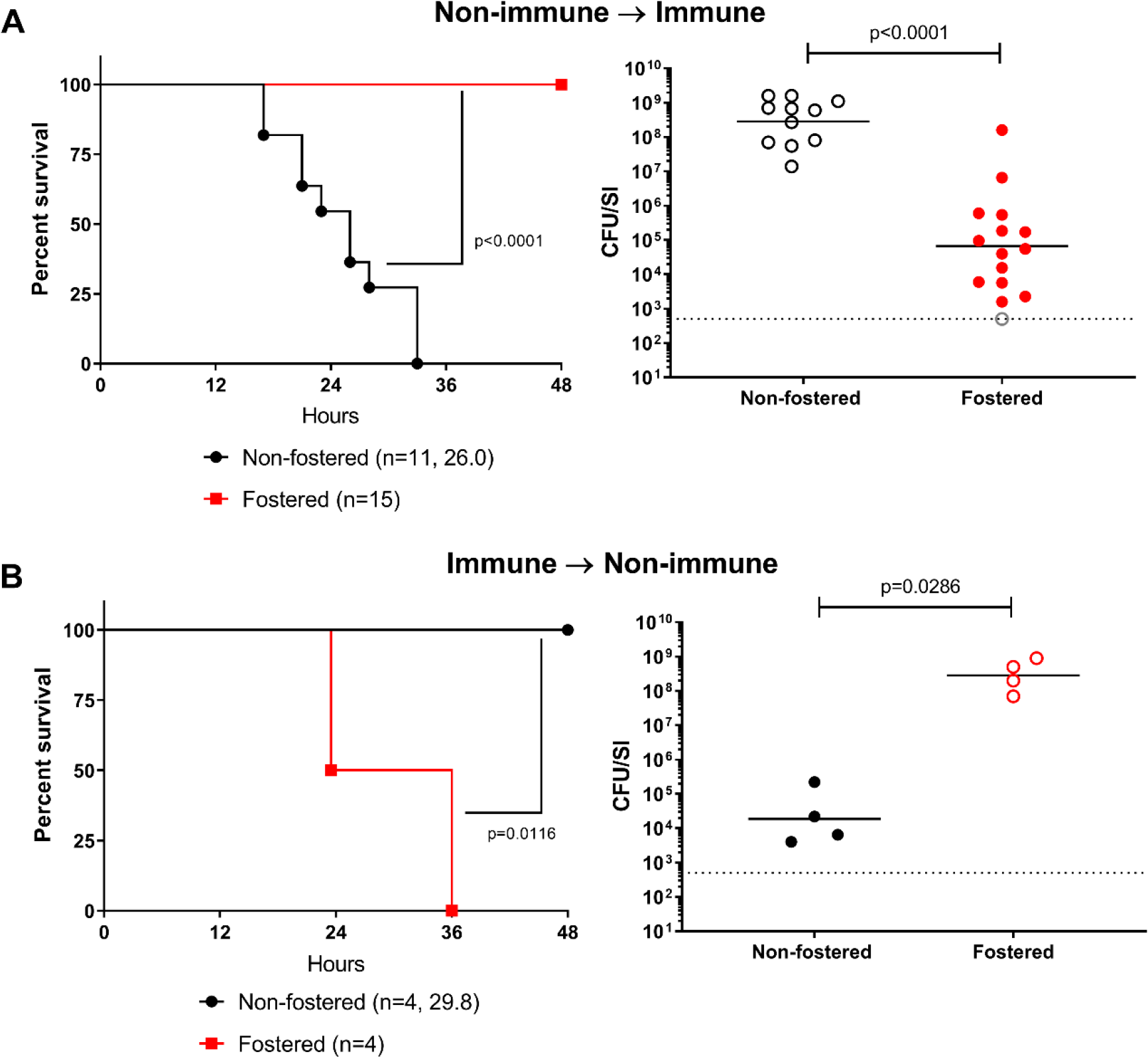
Survival of pups cross-fostered by HaitiV-immunized or non-immunized dams after challenge with HaitiWT. (A) Survival and *V. cholerae* intestinal colonization in pups born to non-immunized dams fostered by HaitiV-immunized dams. (B) Survival and *V. cholerae* intestinal colonization in pups born to HaitiV-immunized dams and fostered by non-immunized dams. Panel labeling, and statistics are as described in Figure 6.

### A single oral dose of HaitiV is sufficient to elicit protective immunity

Finally, to investigate whether the multiple dose regimen was necessary for stimulating protective immunity, we tested whether a single oral dose of HaitiV stimulated protective immune responses in female GF C57BL/6 mice (n=4). Like our studies with the multiple dose regimen, a single dose of HaitiV led to sustained colonization in the mice (Fig S2). HaitiV induced vibriocidal antibody titers comparable in magnitude to those from serially immunized mice (Fig 2C). Litters from singly-immunized mice were also completely protected from disease resulting from HaitiWT challenge, phenocopying pups from the first cohorts (Fig S2).

## Discussion

Vaccines for cholera are being increasingly embraced as public health tools for prevention of endemic cholera and limiting the spread of cholera epidemics [17]. Killed OCVs are efficacious in endemic populations, but live OCVs promise to be more potent, particularly in young children [19]. Here, we showed that the live OCV candidate HaitiV induces vibriocidal antibodies and other immunological correlates of protection against cholera in GF mice and leads to protection against disease in their offspring. Protection in this model was dependent on passively acquired factors in the milk of immunized dams and not transmission or colonization of HaitiV. Although our relatively small cohort sizes precluded rigorous statistical comparisons of immune responses in the immunized mice, oral administration of even a single dose of HaitiV elicited detectable vibriocidal antibodies in all animals. These observations provide strong data establishing HaitiV’s immunogenicity. Additionally, the comparable vibriocidal titers elicited by HaitiV and CVD-103HgR, a live OCV licensed by the FDA for travelers, bodes well for HaitiV’s immunogenicity in humans. Combining the immunogenicity data presented here with our finding that HaitiV can protect from cholera prior to the induction of adaptive immunity [29], suggests that HaitiV may function as a dual-acting agent, providing both rapid-onset short-term protection from disease while eliciting stable and long-lasting immunity against cholera.

Data from the challenge experiments (Figure 6) are consistent with the prevailing notion that serogroup, and to a lesser extent serotype are major determinants of protection against *V. cholerae* challenge [39,46]. Although it is thought that exposure to Inaba strains is more cross-protective than exposure to Ogawa strains, the relative potency of Inaba versus Ogawa vaccines in eliciting dual protection against both O1 serotypes requires further definition, as it has been suggested that both Ogawa and Inaba vaccine strains are good candidates for development [4–7]. A mixture of Ogawa and Inaba serotypes either as distinct strains or one bivalent strain (serotype Hikojima) may be beneficial in broadening the breadth of the immune response to HaitiV [47,48].

The modest protection that HaitiV immunization provided against *V. cholerae* O139 was unexpected. The epidemiology of the original O139 outbreak and experimental studies in rabbits demonstrate a lack of cross-protection between the two serogroups [14,15,39]. Notably, although pups born to HaitiV-immunized dams and challenged with MO10 survived longer than pups born to non-immunized dams, there was little difference in the MO10 intestinal colonization between these groups (Fig 6C). The discrepancy between clinical protection and relatively robust colonization suggests that HaitiV stimulates immune responses to *V. cholerae* factors, like CT, that may contribute to disease but not directly to colonization. The capacity of live OCVs to induce immune responses to in vivo-expressed antigens, including CtxB, is a property that heightens the appeal of live vs killed OCVs [27,49].

Although GF mice enabled us to test the protective efficacy of a candidate live OCV, the absence of the microbiota and resulting improper immune development in these mice, are important caveats to consider. The GF model does not recapitulate the competitive microbial environment that live OCVs will encounter in the human host. We observed similar prolonged shedding patterns for both CVD-103HgR and HaitiV in the GF mice (Fig 1), yet CVD-103HgR is known to be shed by human volunteers at a low frequency for a short period [25,50]. Thus, our findings likely overestimate HaitiV’s capacity to colonize the human intestine. The observation that exposure of HaitiV-immunized dams to SPF-derived pups during the cross-fostering experiments led to the elimination of detectable HaitiV in feces supports the prediction that this vaccine will not stably colonize humans. It is an open question whether transient exposure of naïve mice to HaitiV will also stimulate protective immunity, as has been shown in the context of vaccination with *V. cholerae* outer membrane vesicles [51,52]. The streptomycin-treated mouse model of *V. cholerae* colonization, which allows for temporary intestinal colonization, may also be useful to investigate the duration of colonization required for immunity [53]. Ultimately, the capacity of HaitiV to colonize the intestine and the relationship between colonization and protective immunity will need to be defined in human volunteers.

The immunogenicity of live OCVs in mice has only been investigated in GF animals because adult mice with intact microbiota are refractory to intestinal *V. cholerae* replication and colonization. However, previous studies of live OCVs in GF mice only analyzed immune correlates of protection and not protection against challenge [36,37]. The combination of the neonatal survival assay with the oral GF vaccination model builds on existing knowledge of these mice to assay both the immunogenicity and protective efficacy of live OCV candidates [38]. This model may be a useful addition to existing approaches that probe the molecular bases of vaccine-mediated mucosal protection against pathogens, a topic with significant translational potential that remains poorly understood [52,54]. A recent report employing a similar maternal-infant transmission model in the context of intraperitoneally-delivered heat-killed *Citrobacter rodentium* highlights the versatility of assessing vaccine protective efficacy using the infant progeny of immunized animals as readouts [55]. The broad availability of genetically engineered mice and the relative ease of GF-derivation provides a powerful opportunity to leverage both host and bacterial genetics to explore how live-OCVs can be optimized to better defend against this ancient pathogen.

## Acknowledgements

We thank members of the MKW laboratory for helpful discussions.

## Author Contributions

Conceptualization: BS, TZ, ETR, MKW. Methodology: BS, TZ, ETR, MKW. Investigation: BS, TZ, BF, AA, RB. Supervision: ETR, MKW. Visualization: BS. Writing (Original Draft Preparation): BS and MKW. Writing (Review and Editing): BS, TZ, BF, AA, RB, ETR, MKW.

## Supporting information

**S1 Fig. Duration of protection from HaitiWT in progeny from HaitiV-immunized dams.** Litters were inoculated identically to initial challenge studies and allowed to age normally for up to 14 days (336 hours) before enumeration of intestinal burdens.

**S2 Fig. Effectiveness of a single-dose HaitiV vaccination regimen in C57BL/6 mice.** (A): Fecal shedding of HaitiV from mice given a single dose of HaitiV at Day 0. (B): Vibriocidal antibody titers from singly-immunized mice against either Ogawa (blue) or Inaba (green) target strains. (C): Survival (left) and intestinal colonization (right) of pups from singly-immunized dams challenged with a lethal dose of HaitiWT. The dotted line marks the limit of detection.

**S1 Table.** Bacterial strains used in this study. ET = El Tor, AET = Altered El Tor, Sm = streptomycin, SXT = sulfamethoxazole/trimethoprim, R = resistant, S = sensitive.

| Strain | Notes | Source |
| --- | --- | --- |
| *E. coli* SM10pir | Donor strain for *ctxAB* deletion | [29] |
| *V. cholerae* HaitiWT | O1 Ogawa AET, SmR SXTR, *lac*+ | [10] |
| *V. cholerae* Haiti Δ*ctxAB* | O1 Ogawa AET, SmR SXTR, *lac*+ | This study |
| *V. cholerae* HaitiV | O1 Ogawa AET, SmR SXTS, *lac*- *recA*+ | [29] |
| *V. cholerae* CVD103-HgR | O1 Inaba Classical, SmR, *lac*+ | This study |
| *V. cholerae* N16961 | O1 Inaba ET, SmR, *lac*+ | [56] |
| *V. cholerae* MO10 | O139, SXTR, *lac*+ | [57] |
| *V. cholerae* PIC018 | O1 Inaba ET | [34] |
| *V. cholerae* PIC158 | O1 Ogawa ET | [34] |

**S2 Table.** Multiple comparisons testing of antibody titers in cohort 2. P-values shown were calculated from a Dunnett’s multiple-comparison test comparing Day 14, 28 or 42 mean titers to the mean titer at Day 7. P-values are shown to three significant figures and values < 0.05 are bolded.

| Sample | Day 14 vs. Day 7 | Day 28 vs. Day 7 | Day 42 vs. Day 7 |
| --- | --- | --- | --- |
| Inaba OSP IgA | 0.560 | 0.112 | **0.0197** |
| Inaba OSP IgG | 0.999 | 0.117 | 0.0579 |
| Inaba OSP IgM | **0.0157** | **0.0428** | **0.0357** |
| Ogawa OSP IgA | 0.531 | 0.0828 | 0.146 |
| Ogawa OSP IgG | 0.550 | **0.0205** | **0.0172** |
| Ogawa OSP IgM | 0.0630 | **0.0253** | **0.0343** |
| CtxB IgA | 0.750 | 0.571 | 0.207 |
| CtxB IgG | 0.694 | 0.263 | 0.112 |
| CtxB IgM | 0.267 | **0.0498** | **0.013** |

