## Supplementary figures and images for "Oral immunization with a probiotic cholera vaccine induces broad protective immunity against *Vibrio cholerae* colonization and disease in mice"

### Supplementary Figure 1

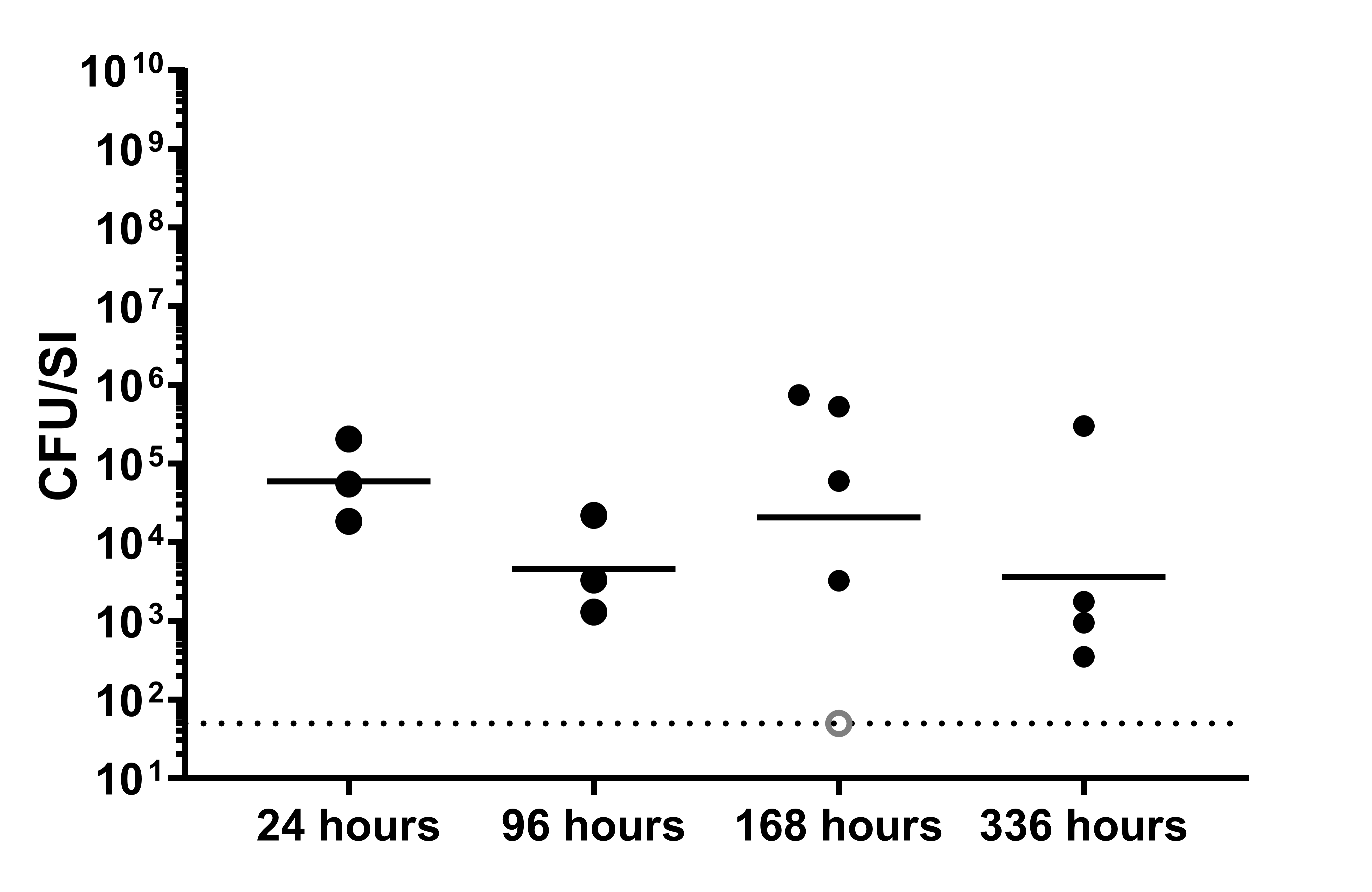

### Supplementary Figure 2

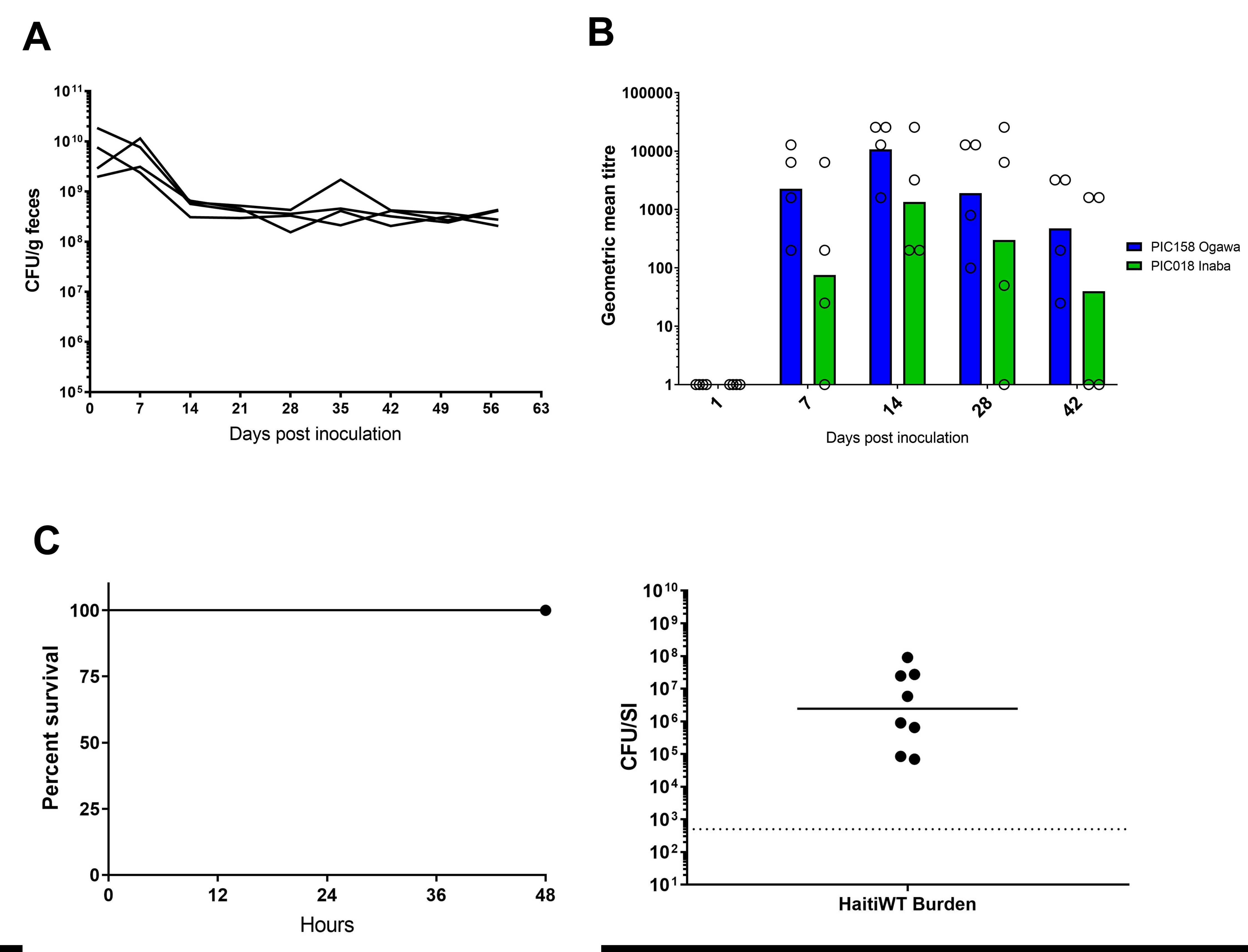
